# Dicarbonyl stress debilitates mesothelial defense against metastasizing ovarian cancer

**DOI:** 10.1101/2024.06.18.599500

**Authors:** Satyarthi Mishra, Kottpalli Vidhipriya, Anchita Gopikrishnan, Sudiksha Mishra, CVS Prasanna, Prosenjit Sen, Ramray Bhat

## Abstract

Chronic metabolic disorders and aging result in accumulation of active dicarbonyls that glycate biomolecules rendering them dysfunctional. Although metabolic aberrations are known to be epidemiologically associated with faster cancer progression, cell biological determinants of such associations remain elusive. The formation of micro-metastases in epithelial ovarian cancer involves its colonization of visceral peritonea through clearance of mesothelia that line the coelom. In this study, we observe that cocultures of immortalized human coelomic MeT-5A mesothelia with human ovarian cancer cells OVCAR-3 and SK-OV-3 show greater infiltration by the latter when exposed to increasing concentrations of the dicarbonyl methylglyoxal (MG). Treatment with increasing concentrations of MG caused death and senescence within human and murine serosal mesothelia. Cells showed higher levels of advanced glycation end products, dysregulated occludens junction protein ZO-1, and disrupted localization of cortical filamentous actin and its regulator ezrin, indicating poor inter-cell adhesion. Time lapse imaging also showed impaired migration for MG-treated single mesothelia and for their collective monolayers. Agent-based computer modeling of coculture dynamics predicted that a combined effect of confluence and migration allows inter-adherent mesothelia to contain the spread of colonizing cancer cells, which was confirmed through coculture time lapses of cancer colonization within higher and lower mesothelial densities. We found ovarian cancer cells showed higher levels of glyoxalase-1 (GLO-1) enzyme, which catabolizes MG, suggesting how they escaped its cytotoxic effects. Consistent with this, treatment of OVCAR-3 with MG concurrently with pharmacological inhibition of GLO-1 showed greater cell death. Our results suggest dicarbonyl stress helps colonizing cancer cells overcome the resistance of natural homoeostatic barriers and its inhibition may, in supplementation with chemotherapy, stem metastasis.

## Introduction

Ovarian cancer is the most lethal among gynecological malignancies (Lheureux et al., 2019). It is the eighth-most common cause of cancer-related deaths among women and upon diagnosis less than 50% show a survival of 5 years or more (Webb & Jordan, 2017). High mortality rates result from its insidious nature of metastasis: observable symptoms are mostly consequential to carcinomatosis resulting in delayed and inadequate management that fails to limit its spread (Bowtell et al., 2015). The incidence of ovarian cancer is higher after menopause and mortality rates are higher when the disease is diagnosed in older women (Lefur et al., 2014). Tetsche and coworkers have observed that age-associated comorbidities contribute to worsening survival in ovarian cancer patients (Tetsche et al., 2008). Of such comorbidities, several are associated with systemic inflammation, including, but not limited to, diabetes. Associations between incidence of cancer and diabetes have been made since the beginning of the last century (Greenwood & Wood, 1914) and epidemiological studies across countries make a strong case for the latter to play an important role in cancer progression (Suh & Kim, 2019) (Gordon-Dseagu et al., 2014)). The presence of Type I or Type II diabetes mellitus has been shown to exacerbate the relative risk of incidence of cancer (including ovarian) by as much as 10% (Tsilidis et al., 2015).

Notwithstanding the confirmation by focused meta-analyses of the associations between ovarian cancer and diabetes (Lee et al., 2013) (Zhang et al., 2017), a histopathological understanding of how one pathological syndrome exacerbates the other remains ill-understood. Circulatory metabolites, such as glucose and triglycerides that are elevated in diabetes may fuel ovarian cancer cells (Lamkin et al., 2009); (Zeleznik et al., 2020). Epigenetic changes associated with diabetes, including elevated levels of insulin-like growth factors, and inflammatory cytokines, such as IL-6 and TNF-α, are also known to potentiate proliferation and migration of ovarian cancer cells (Gallagher & LeRoith, 2015) (Gallagher & LeRoith, 2011) (Amato et al., 2014) (Yu et al., 2009). mIR-29b which belongs to the family of microRNAs dysregulated in diabetes (Kurtz et al., 2014) and plays an inhibitory role in diabetic nephropathy (Gondaliya et al., 2019) inhibits ovarian cancer progression through downregulation of Akt signaling and pyruvate kinase and hexokinase expression (Teng et al., 2015).

While these studies examine the role of diabetes driven- or associated-molecular alterations on cancer cell traits, they neglect how the same functions may regulate the function of the tissue microenvironment within which carcinomatosis ensues. Broadway and coworkers propose a similar hypothesis, wherein senescence driven by a diabetic context of T lymphocytes may regulate omental colonization of ovarian cancer (Broadway et al., 2021). In this study, we focus on the role of a specific metabolite methylglyoxal (MG) in modulating the interactions between tumor cells and their untransformed microenvironmental counterparts. MG is produced as an unstable byproduct of an excess glycolytic flux; it is a highly reactive dicarbonyl compound that reacts with proteins, lipids, and nucleic acids forming advanced glycation (Amadori) products and rendering these biomolecules dysfunctional (Glomb & Monnier, 1995). MG and the resultant dicarbonyl stress are not just strongly associated with diabetes (Schalkwijk & Stehouwer, 2019) but are thought to be causative to both insulin resistance (Moraru et al., 2018) and its organopathic manifestations (Schalkwijk, 2015), which are coincident with histopathological signs of aging. Our study starts by observing the debilitation of an important histological barrier to ovarian cancer colonization: the coelomic mesothelia by MG, followed by an examination of the cytopathological traits that contribute to such debilitation. We posit that glycation-driven dicarbonyl stress may contribute to peritoneopathy that is permissive to the colonization of ovarian cancer. We also investigate how the cancer cells may escape the damaging effects of dicarbonyl stress. This provides a potential insight into how aging, and diabetes (and their accompanying metabolic changes) are risk factors for ovarian cancer metastasis.

## Materials and Methods

### Maintenance and culture of mammalian cells

SK-OV-3 and OVCAR-3 human ovarian carcinoma cells were received as a kind gift from Dr. Rajan R. Dighe, Indian Institute of Science. Immortalized untransformed human mesothelial MeT-5A cells were purchased from ATCC. SK-OV-3 cells were cultured in McCoy’s 5A medium (AL275A, HiMedia) supplemented with 10% fetal bovine serum (FBS) (10270, Gibco). OVCAR-3 cells were cultured in RPMI (AL162A, HiMedia) supplemented with 20% FBS. MeT-5A cells were cultured in M199 medium (AL014A, HiMedia) supplemented with 10% FBS, 5.053 µg/ml insulin, 0.5 ug/ml hydrocortisone, 2.6 ng/ml sodium selenite, 10 ng/ml hEGF (Human Epidermal Growth Factor), 20 mM HEPES. All cells were incubated in a humidified 37 °C incubator with 5% carbon dioxide. For trypsinization, 0.05% trypsin-EDTA (HiMedia) solution was used. Cells were sub-cultured at a confluence of approximately 80% in a ratio of 1:4.

### Culturing spheroids

3% poly-2-hydroxyethyl methacrylate (poly-HEMA; P3932, Sigma-Aldrich) solution was prepared in 95% absolute ethanol. Tissue culture dishes were coated with optimized volumes of the prepared poly-HEMA solution and allowed to polymerize overnight under sterile conditions at room temperature. The dishes were then treated under ultraviolet radiation overnight. Trypsinized cells were seeded according to the requirement of the experiment, in defined medium. Defined medium contained the base medium used for the cell line, supplemented with 0.5 μg/ml hydrocortisone (Sigma-Aldrich, 33 H0888), 250 ng/ml insulin (Sigma-Aldrich, I6634), 2.6 ng/ml sodium selenite (Sigma-Aldrich, S5261)), 27.3 pg/ml estradiol (Sigma-Aldrich, E2758) and transferrin (Sigma-Aldrich, T3309). Depending on the experimental requirements, the cultures were then incubated in a humidified 37 °C CO_2_ incubator for 24 or 48 h.

### Drug treatments

MeT-5A, OVCAR-3 and SK-OV-3 cells were treated with the dicarbonyl stress-inducing compound methylglyoxal (MG) (M0252, Sigma-Aldrich) for 48-72 h.

OVCAR-3 cells were treated with s-p-Bromobenzylglutathione cyclopentyl diester (Glo-1 inhibitor) (SML1306, Sigma-Aldrich) for 72 h.

### Resazurin assay

A cell-permeable redox indicator resazurin (R7017, Sigma-Aldrich) was used to assess the viability of ovarian cancer cells. Viable cells with active metabolism reduce resazurin to a pink fluorescent compound resorufin, and the amount of resorufin produced is proportional to the number of viable cells. The assay was performed by seeding 3000 cells per well of a 96 well plate. After 12 h, they were treated with different concentrations of MG and incubated for 72 h. Puromycin (1 μg/mL) was used as a positive control for cell death. After the incubation period, 10 μg/μL resazurin was added to each well and incubated for 1.5 h. Fluorescence was measured using a microplate fluorometer (Tecan Infinite M Plex) equipped with a 560 nm excitation/590 nm emission filter set. IC50 values were calculated using a dose-response curve in Prism GraphPad (Arora et al., 2023).

### Wound Healing / Scratch Assay

MeT-5A were seeded in a 24 well plates and let to form a confluent monolayer after which a sterile pipette tip was used to make scratch. The cells were treated with 50 µM and 100 µM of MG and the wound healing was imaged every 24 h for 2 days. The quantification was done according to the formula (Rötzer et al., 2016).

### Mesothelial clearance assay; Co-culture setup

Either one or both cells (cancer cells and mesothelial cells) in the co-culture setup were chosen as fluorescent to be able to lineage trace them once the cells have intermixed. The cells were made fluorescent by stably over-expressing fluorescent proteins (green fluorescent protein (GFP) / red fluorescent protein (RFP)) in the cells using a lentiviral transduction approach. The transduced cells are selected with puromycin and enriched by fluorescence assisted cell sorting (FACS). For the assay, trypsinized MeT-5A cells were seeded in eight-well chambered cover glasses at a density of 40,000 cells/well. The cells were allowed to form monolayers of mesothelial cells for 36-48 h in a humidified 37°C CO_2_ incubator in absence and presence of methylglyoxal (50 & 100 µM). Simultaneously, trypsinized ovarian cancer cells were seeded in poly-HEMA coated 35 mm tissue culture-treated dishes for the formation of spheroids and cultured in a humidified 37 °C CO_2_ incubator. At the end of 36-48 h, the ovarian cancer spheroids were added into the eight-well chambered cover glasses with MeT-5A monolayers. Immediately, epifluorescent timelapse imaging with appropriate settings was set up and started for the duration and time-interval relevant to the experiment. The acquired images were processed and analyzed for area of spread/area of clearance using FIJI ImageJ.

### Virus preparation

HEK293FT cells were purchased from Invitrogen (R70007) and cultured in DMEM High Glucose, supplemented with 10 % FBS and 2 mM glutamine. Trypsinized HEK 293 FT cells were seeded at 50 % confluency in DMEM High Glucose containing 10 % FBS and 2 mM glutamine. After overnight incubation in a humidified 37°C CO_2_ incubator, the medium was replaced with serum free-DMEM High glucose for 1 hour before adding the plasmids in the following ratio (as per Cribbs et al., 2013): psPAX2 and pMD2.G (packaging plasmids, 5000 ng: 2000 ng) gene of interest in pCDH vector: pCDH-tGFP-T2A-Puro or pCDH-RFP-T2A-Puro (5000 ng); pCDH-T2A-Puro was used as vector control. The plasmids were mixed in the above ratio in Turbofect (Invitrogen, USA), incubated at room temperature for 30 min, before adding to the cell monolayer. The cells were incubated with the mixture for 6 h in a humidified 37°C CO_2_ incubator, to allow plasmid uptake. At the end of 6 h, media was changed and DMEM High Glucose supplemented with 10% FBS and 2 mM glutamine was added. Cells were incubated again for 48 h, or 72 h, for the production of viral particles. Medium with lentiviral particles was harvested at 48 h and subsequently at 72 h post-transfection. The harvested medium was centrifuged at 1200 rpm for 5 min and the supernatant was filtered using a 0.2 μm filter. The filtered virus-containing suspension was directly used for transduction or concentrated using Lenti-X concentrator (Takara, 631231) using the manufacturer’s protocol. The leftover viral suspension was stored as single-use aliquots at -80 °C for future usage.

### Viral transduction of MeT-5A, SK-OV-3 and OVCAR-3 cells

Cells were seeded at about 50 % confluency and, after overnight incubation, were treated with complete medium containing Polybrene (Sigma, USA) at a final concentration of 8 μg/ mL and lentiviral particles. The cells were incubated with the mixture for 48 h, in a humidified 37°C CO_2_ incubator. Transduced cells were then selected using complete medium containing 10 μg/ mL Puromycin dihydrochloride (Sigma, USA) for 48 h. The selected cells were trypsinized and maintained in the same puromycin pressure for two passages, following which it was reduced to 5 μg/ mL for the next two passages before withdrawing puromycin. Cells expressing high levels of fluorescent proteins were sorted using flow cytometry (BD Influx cell sorter).

### Ex-vivo murine mesentery dissection

BALB/c female mice (4-6 weeks-old) were used for dissecting out mesenteric tissues with due ethical clearance. Mice were euthanized by cervical dislocation. The abdomen was surgically cut open and their mesentery were strung to the bottom side of transwell inserts (Boyden chambers without the membranes) using a surgical thread. The transwells with strung mesentery were then placed in a sterile 24-well tissue culture plate, and medium was added into the transwell above and below the insert into wells of the plate.

### Live-dead assay using Calcein-AM and propidium iodide

Calcein-AM (for staining live cells, C1430, Invitrogen) was added to the cells at a concentration of 2 μM, directly in culture and incubated for 30 min. After 30 min, cells were then washed and put in a fresh medium to remove AM ester. Propidium iodide (for staining dead cells, BMS500PI, Invitrogen) at a concentration of 50 µg/ml, was then added, incubated for 5 min, and followed by two washes with PBS. The cells were then imaged either by epifluorescence or confocal imaging microscope.

### Immunostaining

Eight-well chambered cover glasses containing cells or mesenteric explants on transwells were fixed using 3.7 % formaldehyde (24005, Thermo Fischer Scientific) at 4 °C for 20 min. After fixation, the cells or explants were taken for further processing or were stored in 1X PBS at 4°C. Cells and explants were permeabilized using 0.5 % Triton X-100 (MB031, HiMedia) in PBS, at room temperature for 2 h. Blocking was achieved using 3 % BSA in PBS containing 0.1% Triton X-100. The solution was added to the cells and incubated at room temperature for 1 hour. The cultures were then incubated in diluted (1:500) primary antibodies at 4 °C overnight. This was followed by three washes for 5 min each, using 0.1 % Triton X-100 in PBS. Then secondary antibody was diluted as per manufacturer’s recommendation and added to the cultures. Simultaneously, the cultures were counterstained with phalloidin for F-actin visualization. This was incubated in the dark, at room temperature for 2 h. DAPI (D1306, Thermo Fischer Scientific) was added at a dilution of 1:1000 and incubated in darkness for 10 min. The cultures were washed thrice with PBS containing 0.1 % Triton X-100 for 5 min each. Negative controls were included in each experiment (no primary antibody was added). The list of catalogue numbers for the primary antibodies, phalloidin and DAPI used in immunostaining follows: Alexa Fluor™ 488 Phalloidin (A12379, Thermo Fischer Scientific), Alexa Fluor™ 568 Phalloidin (A12380, Thermo Fisher Scientific), Anti-Carboxymethyl Lysine antibody [CML26] (ab125145, abcam), Ezrin Polyclonal Antibody (PA5-82769, Thermo Fischer Scientific), Anti-ZO1 tight junction protein antibody (ab96587, abcam), DAPI (D1306, Thermo Fisher Scientific)

### Microscopic image acquisition and processing

#### a. Timelapse imaging

Experiment-specific cultures with either cells or spheroids cultured on ECM scaffolds or on mesothelial cells in eight-well chambered cover glasses, were placed in the Tokai stage-top incubator of an Olympus IX73 microscope. The settings for the incubator were maintained at 37 °C, with 5 % CO2 and sterile water being added in the designated space for humidification. The cellSens imaging software was turned on along with all the connected parts associated with imaging, including the Orca Flash LT plus camera (Hamamatsu). The laser settings were made depending on the channels used, and the multipoint timelapse was carried out for 24-48 h, with a time-interval of 10-30 min depending on the experimental requirements.

#### b. Confocal imaging

According to the experiment being performed, the fixed/live samples were taken and placed on the microscope stage. The imaging was done using an Olympus IX83 inverted fluorescence microscope fitted with Aurox structured illumination spinning disk setup, the Leica SP8 confocal microscope and Andor Dragonfly (CR-DFLY-502) confocal imaging platform. The settings were made according to the experiment, with appropriate controls. The images acquired via the Leica microscope were processed on the LASX software.

### Imaging analysis/quantification

#### a. Area of spread

Using FIJI ImageJ (Schindelin et al., 2012), area of spread of spheroids was measured in pixel squares. At an initial time point, the area was measured and called as initial area (A). At the end-time point the area was measured and called as final area (B). The fold change in spreading area was calculated by the formula (B/A).

#### b. Measurement of CML expression data

The levels of CML expression in control and MG-treated mesothelia were measured from the immunostaining signal using the CellProfiler software (Stirling et al., 2021).

#### c. F-actin plot profile

The expression of filamentous actin (F-actin) across the span of the cell in control vs MG-treated mesothelia was measured from the Phalloidin staining using the Plot Profile plugin of ImageJ (Schindelin et al., 2012).

#### d. Distance measurements

Accumulated distances travelled and Euclidian distances traversed by single cells were measured from the phase contrast videography using MTrackJ plugin (Meijering et al., 2012)on FIJI ImageJ. Single cells were tracked manually for 4-6 h of time-lapse videography.

### Compucell 3D model for the clearance of senescent mesothelia

Compucell3D (CC3D), the computational simulation framework is derived from the Cellular Potts Model (CPM) or the Glazier-Graner-Hogeweg (GGH) model. The model has been employed to simulate the movement of cells through biologically complex milieu. The CPM is a lattice-based discrete model in which movement in space and time is simulated through an energy minimizing procedure (Graner & Glazier, 1992; Swat et al., 2012). Each cell consists of several lattice sites (pixels). Each configuration is associated with an effective energy, or Hamiltonian (H), which is calculated based on properties such as volume, surface area, contact energies, or external properties. Time evolution is simulated through Monte Carlo Steps (MCS) which involve random changes of lattice site occupations and the changes that decrease the energy are more likely to occur. The Hamiltonian of the system has been described in Thapa et al., 2023.

In this paper, a simple two-dimensional model 140*140*1 using a square pixel lattice was used. The simulation time was set to 10000 MCS as per the corresponding normalized length of the experimental time-lapse videos. The lattice is composed of the medium, mesothelial cells (NON_SEN), ovarian cancer cells (CANCER), and a frozen wall that mimics a closed system such as enclosed peritoneal cavity. This control model has contact energies of cancer and mesothelia to be 50 and 10 reflecting higher inter adhesion strength of mesothelia). Impairment of motility signifies decreasing the motility value to zero. Impairment of mesothelial adhesivity implies an increase in contact energy to resemble epithelial inter-adhesive strength. Division is rates for mesothelia are decreased to three fourths current rates. To compare the simulation time to the timescale of senescent mesothelial clearance by ovarian cancer cells, a calibration was performed. The division time of ovarian cancer cells in experiments (SK-OV-3 cells) was 18 h and the division time of ovarian cancer cells in simulations was 750 MCS. This factor was used to calibrate the simulations time to real-time and the comparison between the computational model and senescent mesothelial clearance experiments was made.

### Statistics

All graphs are plotted, and statistical analyses were performed using GraphPad Prism version 10.2.3 for Mac OS X, GraphPad Software, Boston, Massachusetts USA, www.graphpad.com. Statistical significance was calculated using either paired or unpaired Student’s t-test with Welch’s correction or One-way ANOVA with Dunnett’s post hoc multiple comparison. All experiments were performed three times unless specified.

## Results

### MG exposure enhances the colonization by ovarian cancer spheroids on untransformed mesothelial monolayers

To mimic peritoneal colonization of epithelial ovarian cancer (EOC) spheroids, untransformed mesothelial cells MeT-5A expressing RFP were seeded as monolayers. Such layers were then cultivated under control serum-supplemented conditions or with 50 µM or 100 µM methylglyoxal (MG). The concentration we chose to test was based on preliminary screens where we observed insignificant differences in cellular behavior at concentrations lower than 50 µM with immortalized cell lines. MG levels within cells have been estimated to be between 120 nm to 400 µM (Beisswenger et al., 1999; Lapolla et al., 2005; Scheijen & Schalkwijk, 2014). Kong and colleagues have recently estimated that treatment of cells with 100-500 µM MG leads to an intracellular concentration of 100-700 nM, indicating our treatments are within physiological relevance (Kong et al., 2024). In parallel, ovarian cancer cell lines, OVCAR-3/SK-OV-3 expressing GFP were cultured on low adhesion substrata so as to allow them to form spheroids (Langthasa et al., 2021) and after 48 h, added to MG-treated mesothelia. Medium was replenished with MG every 24 h to maintain the dicarbonyl stress and the cocultures were subjected to time lapse imaging for 24-48 h (Figure 1A). At the end of 48 h, we observed solitary cancer spheroids or single cancer cells (OVCAR-3, Figure 1B) and (SK-OV-3, Figure 1C); green fluorescence) in control monolayers (left). In contrast progressively larger areas of the mesothelial monolayer space (red fluorescence) was occupied by cancer cells as MG concentrations were increased (Figure 1B and C, middle and right).

**Figure 1.**
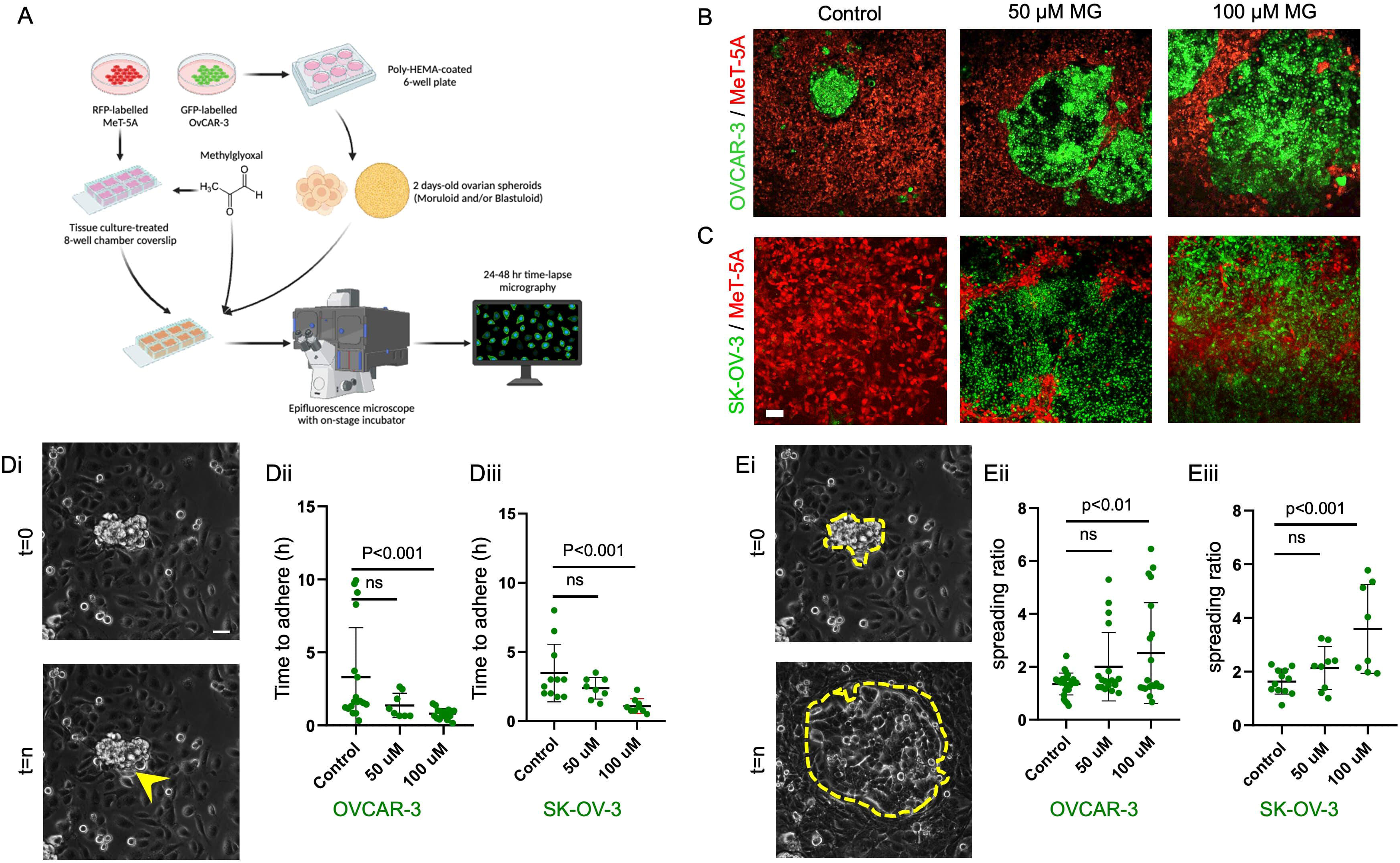
Ovarian cancer spheroid adhere and spread faster on MG-treated mesothelial monolayers. (A) Schematic representation of the experimental design adopted for investigating the mesothelial clearance in a coculture of cancer cells and untransformed control and methylglyoxal (MG)-treated mesothelia expressing distinct fluorescent reporter proteins and imaged using epifluorescence microscopy. (B, C) Laser confocal micrographs of cocultures consisting of OVCAR-3 (B) and SK-OV-3 (C) both expressing GFP within a confluent patch of mesothelia (RFP-expressing MeT-5A) cultured as control (left) and with 50 µM (middle) and 100 µM (right) MG. (D) Brightfield micrographs of spheroid-mesothelial cocultures at the start (top) and end (bottom) of the examined time lapse with yellow arrowhead showing the first instance of spheroidal adhesion through a flattened cell protrusion (Di). Graphs showing time to adhesion of OVCAR-3 (Dii) and SK-OV-3 spheroids (Diii) on control and MG-treated MeT-5A monolayers. (E) Brightfield micrographs of spheroid-mesothelial cocultures at the start (top) and end (bottom) of the examined time lapse with yellow dotted lines showing the spread spheroidal contour at the start and end of time lapse (Ei). Graphs showing spreading ratio of OVCAR-3 (Eii) and SK-OV-3 spheroids (Eiii) on control and MG-treated MeT-5A monolayers. Scale bars for B, C: 100 µM and D, E: 20 µM. Quantification done using one way ANOVA with Tukey’s post hoc comparison. Schematic made using BioRender. Experiments performed n=3 times.

To quantify these observations in real time, we estimated the time required for the cancer spheroids to adhere within the mesothelial monolayers by following the time lapse, and detecting the first substratum-adherent cell that could be visualized as exiting the spheroids (Figure 1Di; images taken at the beginning and specific time points of the time lapse). For OVCAR-3 spheroids, the mean time to adhere on control mesothelia (3.3h) was significantly higher than on 50 µM or 100 µM treated mesothelia (1.3 and 0.8 h respectively; Figure 1Dii; p<0.001 estimated using one way ANOVA with Tukey’s post hoc comparison). Similarly, for SK-OV-3 spheroids, the mean time to adhere on control mesothelia (3.4h) was significantly higher than on 50 µM and 100 µM treated mesothelia (2.3 and 1 h respectively; Figure 1Diii; p<0.001 estimated using one way ANOVA with Tukey’s post hoc comparison). Importantly, the ratio of final spread area of the spheroids to the initial spheroid size i.e. quantitative estimate of colonization (Figure 1Ei), for OVCAR-3 in case of 50 µM and 100 µM treated mesothelia was 2.0- and 2.5-fold compared with on control mesothelia (1.3-fold) (Figure 1Eii; p<0.001 estimated using one way ANOVA with Tukey’s post hoc comparison). For SK-OV-3 spheroids, the mean spreading ratio in case of 50 µM and 100 µM treated mesothelia was 2.1- and 3.6-fold compared with on control mesothelia (1.3-fold) (Figure 1Eiii; p<0.001 estimated using one way ANOVA with Tukey’s post hoc comparison). The increase in colonization of cancer cells amidst mesothelia in the presence of MG can be interpreted by their direct potentiation of cancer cells or/and a debilitation in the physiology of mesothelial layers. We began by testing the latter first.

### MG glycates mesothelia, alters cytoskeleton organization and induces cell death

A highly reactive dicarbonyl, MG reacts non-enzymatically with proteins and forms advanced glycation end products (AGEs) at a rate manifold higher than glucose, thereby making it one of the most potent glycating agents implicated in diabetes like metabolic disorders (N. Ahmed, 2005; Allaman et al., 2015). Typically, MG reacts with the lysine and arginine residues to predominantly form AGE adducts called N-epsilon-(carboxyethyl)lysine (CML) and its homolog N-epsilon-(carboxymethyl)lysine (CML) (M. U. Ahmed et al., 1997; Teerlink et al., 2004) (Figure 2A). CML or carboxymethyl lysine is formed by the non-enzymatic Schiff base reaction of glucose with proteins, followed by an Amadori rearrangement and oxidation that leaves only a carboxymethyl group attached to the lysine. The levels of CML adducts accumulate over time and have been used as an indicator of both serum glucose levels and oxidative protein damage. As observed from immunocytochemical staining, levels of CML were significantly higher in MeT-5A mesothelia exposed to increasing concentrations of MG compared with control conditions (Figure 2B; CML fluorescent signals shown in green) with the integrated CML intensity significantly higher in MG treated cells (p<0.01 and <0.001 for 50 µM and 100 µM, respectively; estimated using one way ANOVA with Tukey’s post hoc comparison). The altered shapes of cells in the previous assay motivated us to investigate the nature of cellular filamentous actin using phalloidin (green), which showed an ectopic diffuse localization when mesothelia were treated with 50 µM and 100 µM of MG (Figure 2C with schematic depiction of the representative localization shown in Figure 2D). Representative traces of the phalloidin signals across single control and MMG-treated cells showed cortically localized F-actin in control mesothelia with a progressive de-corticalized diffusive localization upon MG treatment (Figure 2E). The significant difference in cortical relative to cytoplasmic signals of F-actin in control cells became insignificant upon MG treatment (Figure 2F; p<0.001 for control; estimated using one way ANOVA with Tukey’s post hoc comparison). Using a resazurin assay, we measured and observed a potential decrease in viability of MeT-5A cells when exposed to increasing concentrations of MG (Figure 2G; IC_50_ concentration measured to be 133 µM). This was further confirmed by increased nuclear localization of propidium iodide, a DNA intercalating dye which is impermeable to living cells (Figure 2H; green signals due to Calcein AM). To validate our findings, peritoneal explants from 6-week C57Bl/6 mice were dissected and cultured ex vivo under exposure with MG (schematic depiction shown in Figure 2I): greater PI and lower Calcein AM signals were observed respectively in 50 µM and 100 µM of MG-treated cells compared with controls (Figure 2J). The altered spatial localization and the disruption in mesothelial apposition led us to investigate if MG disrupts localization of cytoskeletal regulators.

**Figure 2.**
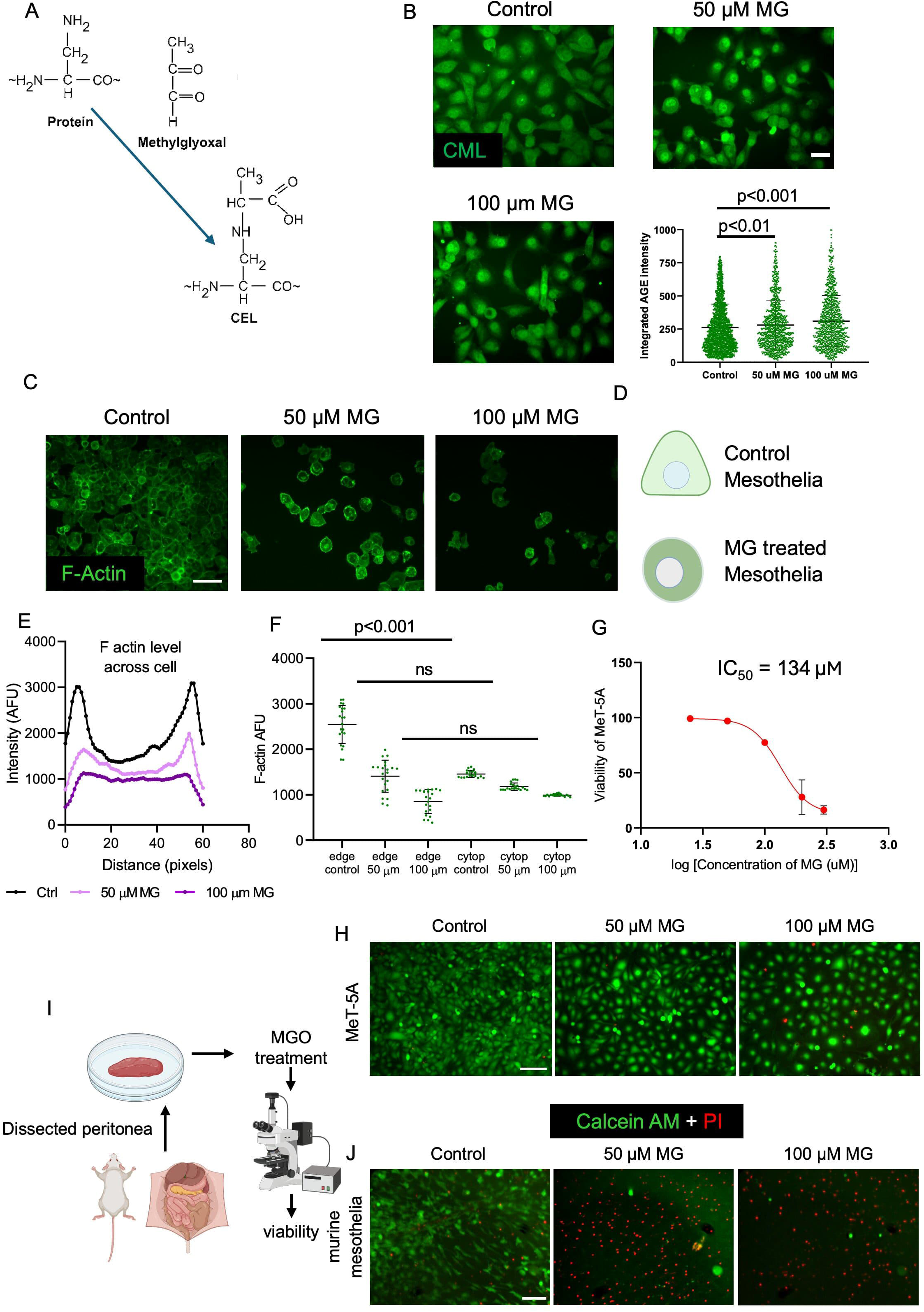
MG-treated mesothelial monolayers show increased AGE levels, diffused/decorticalized filamentous actin and decreased viability. (A) Chemical reaction depicting the conjugation of methylglyoxal with proteins to form N-epsilon-(carboxymethyl)lysine (CML), a major advanced glycation end-product (AGE) implicated in Type 2 diabetes. (B) Confocal fluorescent micrographs of CML immunostaining in control and MG-treated mesothelial cells. Graph plotting the levels of CML formation (each dot representing an individual cell) in control and MG-treated mesothelial cells measured from CML immunostaining signals. (C) Confocal fluorescent micrographs of F-actin staining (with Phalloidin) in control and MG-treated mesothelial monolayers. (D) Schematic representation of a mesothelial cell shape in control and MG-treated conditions viz. cortical distribution of F-actin in control vs diffused F-actin in MG-treated mesothelia. (E) Graph plotting F-actin level across the span of a cell, from one end to the other, measured using Phalloidin staining signals as traces in control (black), 50 µM MG and 100 µM MG treated mesothelia. (F) graph showing mean level of F-actin at the edge and in the cytoplasm of control (black), 50 µM MG and 100 µM MG treated mesothelia. (G) Graph plotting viability of untransformed mesothelia cultured increasing concentrations of MG (IC_50_ = 134 µM). (H) Confocal fluorescent micrographs of MeT-5A mesothelial monolayers stained with Calcein AM (green; live) and Propidium Iodide (red; dead) depicting the viability in control cells and upon 50 µM and 100 µM MG treatment. (I) Schematic representation of the steps involved in the isolation of and subsequent experimentation with *ex vivo* murine mesentery. (J) Fluorescent micrographs of *ex vivo* murine mesenteries stained with Calcein AM (green; live) and Propidium Iodide (red; dead) depicting the viability in control cells and upon 50 µM and 100 µM MG treatment. Scale bars for B, C, D & E: 100 µM: 20 µM. Quantification done using one way ANOVA with Tukey’s post hoc comparison. Schematic made using BioRender. Experiments performed n=3 times.

### Methylglyoxal disrupts epithelial homeostasis

Mesothelia act as active cellular barriers within coelomic cavities surveiling the serosal surfaces against potential pathological insults (Mutsaers et al., 2016). Thus, although they may be tightly apposed to each other using cellular junctions (Konings et al., 1989; Levai et al., 2023), their motility- and permeability-based functions renders makes such junctions more dynamic, allowing tight junctional proteins to reside both at cell surfaces and in cytoplasm (Retana et al., 2015).Therefore, unsurprisingly immunocytochemical detection of Zonula occludens-1 (ZO-1 also known as Tight junction protein-1) showed them to localize in the cell surfaces as well as cytoplasm in control mesothelia. ZO-1 is a 220-kDa peripheral membrane protein connecting tight junction complexes with the underlying actin cytoskeleton (Figure 3Ai shows schematic depiction of ZO-1 localization, see also Figure 3Aii). Treatment with MG led to an increasingly punctate perinuclear localization suggesting an impairment in tight junctions with MG-treated monolayers (Figure 3Aiii and iv right; confocal micrographs of MeT-5A cells stained for ZO-1 (green) and DNA (blue)).

**Figure 3.**
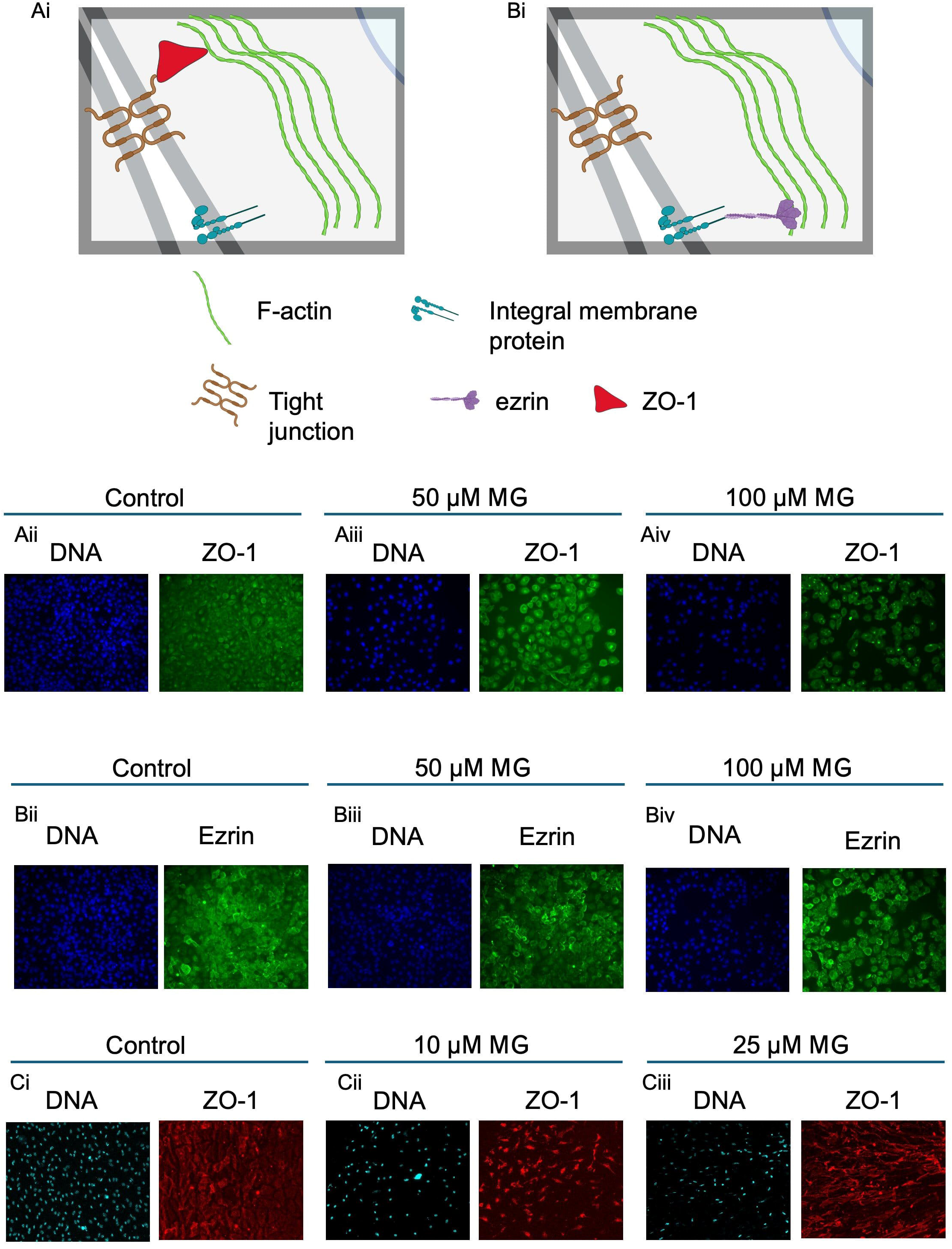
MG-treated mesothelial monolayers show disrupted localization of ZO-1 and ezrin. (Ai) Schematic representation of how Zonula adherens-1 (ZO-1) anchors the tight junctions to the actin cytoskeleton (Bi) Schematic representation of how Ezrin anchors integral membrane proteins to the underlying actin cytoskeleton (Aii-iv) Confocal fluorescent micrographs of control and 50 µM MG and 100 µM MG-treated mesothelia stained for DNA (blue, using DAPI), and for ZO-1 (green, using cognate antibody) (Bii-iv) Confocal fluorescent micrographs of control and 50 µM MG and 100 µM MG-treated mesothelia stained for DNA (nucleus, blue, using DAPI), and for ezrin (green, using cognate antibody). (Ci-iii) Confocal fluorescent micrographs of control and 50 µM MG and 100 µM MG-treated murine mesenteries stained for DNA (blue, using DAPI), and for ZO-1 (red, using cognate antibody) Scale bars for Aii-iv and Bii-iv: 100 µM. Schematic made using BioRender. Experiments performed n=3 times.

Using immunocytochemistry, we also investigated the localization of Ezrin (also known as cytovillin or villin-2), a cortically located protein that crosslinks the actin cytoskeleton and integral membrane proteins (Casaletto et al., 2011). Ezrin has been recently implicated in the regulation of cell size and intercellular tension with its levels regulating the propensity for fracturing in epithelial monolayers (Chouhan et al., 2024; Rouven Brückner et al., 2015) (Figure 3Bi shows schematic depiction of Ezrin localization, see also Figure 3Bii). We observed that treatment with increasing levels of MG resulted in a decorticalization and a diffuse cytoplasmic distribution of Ezrin through the mesothelial soma (Figure 3B iii and iv; confocal micrographs of MeT-5A cells stained for Ezrin (green) and DNA (blue)).

In addition to human mesothelia, we also observed ectopic localization of ZO-1 within murine mesenteries that were exposed to relatively lower concentrations of MG (10 µM and 25 µM) along with a depletion of cells (as observed through the number of nuclei in Figure 3Ci-iii; DNA blue ZO-1 red). Murine mesenteries did not stain clearly for Ezrin, which could be explained by non-reactivity of the antibody across species.

### Methylglyoxal inhibits dispersed and collective migration

The alterations in actin cytoskeletal pattern as well as that of its regulators such as ZO-1 and Ezrin led us to investigate whether dicarbonyl stress also affected mesothelial migration. Experiments measuring both collective motility and single cell migration of mesothelia were performed. For measuring collective cell motility, scratches were made in a high-density mesothelial monolayer and were observed to study their closure as a function of time in controls and under 50 µM and 100 µM MG treatment (Figure 4A). While the collectively motile mesothelia filled the scratch within 3 days, treatment with 50 & 100 µM MG inhibited the migration of MeT-5A and impaired the closure of the gap (Figure 4B).

**Figure 4.**
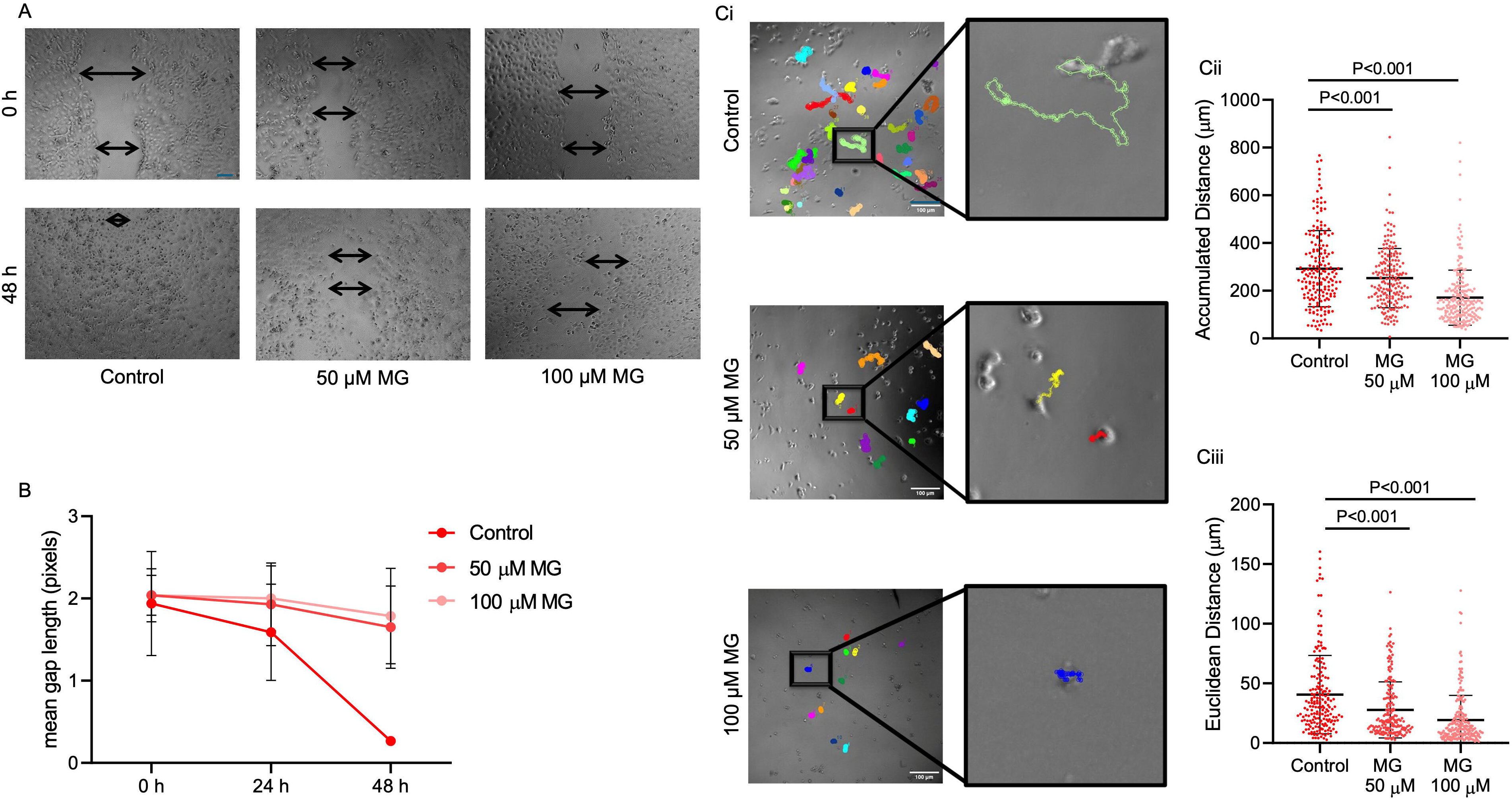
MG-treated mesothelial cells show impaired dispersed and collective migration. (A) Brightfield micrographs of control and 50 µM MG and 100 µM MG-treated mesothelial monolayers at the initial (0 h; top) and later (48 h; bottom) timepoints of the scratch assay with the black arrows showing the leading edge on either side of the gap. (B) Graph plotting the gap measured in control and MG-treated mesothelial monolayers at the beginning of the experiment (0 h), after 24 h and after 48 h. (Ci) Brightfield micrographs of motility trajectories of control and 50 µM MG and 100 µM MG-treated mesothelial cells, with the respective insets depicting the motility trajectory of one cell from each condition. (Cii,iii) Graphs plotting the total distance travelled (ii) and net displacement (iii) of control and 50 µM MG and 100 µM MG-treated mesothelia as measured using track lengths. Scale bars for A & C: 100 µM. Quantification done using one way ANOVA with Tukey’s post hoc comparison. Experiments performed n=3 times. (For reference, check Supplementary Videos S1-S6)

To investigate dispersed or single cell migration, mesothelia were cultivated at low density and treated with MG and imaged under time lapse at intervals of 15-20 mins (Figure 4Ci, Supplementary Videos S1-S6). Mean accumulated distance, i.e. the total distance traversed by a cell over the entire period for control mesothelia were 300 mm, while in case of 50 & 100 µM MG treatments, were found to be 250 and 180 mm respectively (Figure 4Cii; significance measured using one way ANOVA with Tukey’s post hoc comparison). Euclidian distance i.e., the measure of net displacement of the cell from initial to final position, was found to be around 40 mm in control, and 25 and 18 mm under 50 & 100 µM MG respectively (Figure 4Ciii; significance measured using one way ANOVA with Tukey’s post hoc comparison).

### A computational multiscale model predicts the relevance of MG for colonization

In order to decipher which among the cytological effects we observed in mesothelia as a result of MG exposure were instrumental to the cancer colonization coculture seen in Figure 1B, we set up an experimentally motivated multiscale agent-based computational framework based on the Cellular Potts model (Graner & Glazier, 1992; Thapa et al., 2023). We chose to begin our Monte Carlo simulations with two cancer cells (red) and eight mesothelia (green) within a bounded 2D substrata environment. The asymmetry in initial cell number was to model a serosal surface environment, wherein mesothelia were greater in number and could outnumber and surround cancer cells (Figure 5Ai and ii). In addition, the inter-adhesion strength of cancer cells was kept to a fifth strength of that of mesothelia give the tendency of the latter to form apposed monolayer sheets. We ran the simulations for a fixed length of 10000 Monte Carlo steps, wherein to mimic the effects of MG, we decreased the (i) inter-mesothelial adhesion (to capture the effects of tight junctional disruption and spatial disruption of actin) (ii) mesothelial proliferation rate (to mimic the decrease in cell viability) and (iii) mesothelial motility (in association with the decrease in cell migration dynamics), individually, in pairs, and all three traits concurrently (Figure 5Aiii); we then recorded the extent of cancer colonization through a ratio of cancer cell number/mesothelial number at the end of simulations and also compared the spatial distribution of the digital coculture at the end of simulations with our experimental results. Disruption in motility by itself had little effect on the interrelationships between cancer cells and mesothelia compared with control (Figure 5Aiv). In contrast, impairment in division rate of mesothelia allowed a larger population of cancer cells to occupy the digital serosal space. However, an interpenetration of the two cell populations which is evident to a mild extent in OVCAR-3 and especially for SK-OV-3 was not seen in this case where the two digital cell populations remained well sorted (Figure 5Av). While increase in interpopulation penetration and therefore, digital cancer invasion was seen when adhesion was solely disrupted, the extent of colonization measured through the relative population cell number ratio remained close to control (Figure 5Avi). While combined disruption in adhesion and motility separately, did not increase colonization (Figure 5Avii), an impairment of proliferation simultaneously with motility or adhesion allowed digital cancer cells at the end of the simulations to colonize to a greater extent (Figure 5Aviii and ix) although in the former, the invasive spatial phenotype was not evident and in the latter colonization was equal to or lower than that achieved when only division was impaired. (Figure 5B; significance computed using one way ANOVA with Tukey’s post hoc comparison).

**Figure 5.**
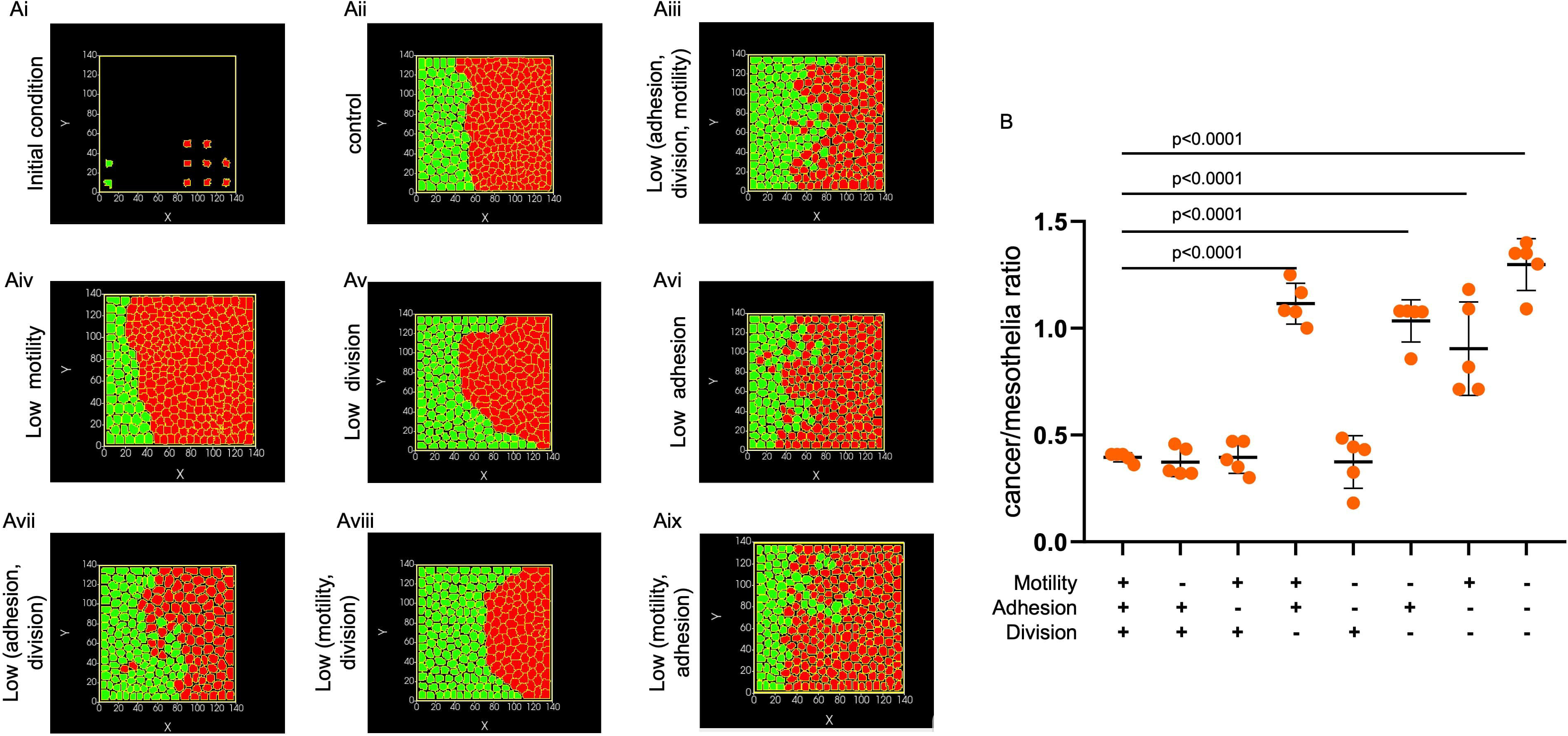
Multiscale simulations predict motility, cell adhesion and viability of mesothelia are vital to their homoeostasis. (A) Simulation images showing (i) initial configuration of red (cancer) and green (mesothelia) at the start of the simulations and (ii) at the end of 10000 Monte Carlo Steps (MCS) for control, and (iii) for the condition when mesothelial motility, division rate, and adhesion are impaired, where (iv) motility, (v) division rate and (vi) adhesion are individually impaired, and where (vii) adhesion and division, (viii) motility and division, and (ix) motility and adhesion are impaired in a pairwise manner. (B) Graph showing the ratio of cancer cell number to mesothelial number for all the conditions in (A). Quantification done using one way ANOVA with Tukey’s post hoc comparison. Simulations performed n=5 times.

The strongest significant results for spatially consistent invasion of cancer cells were observed when all three traits of mesothelia: motility, adhesion and proliferation were impaired. This suggests that whereas proliferation is sufficient to ensure greater cancer colonization, the spatial interfacing we observe in experiments necessitates the relevance of impairment in mesothelial adhesion through effects on cytoskeletal elements. Greater invasion of cancer cells is achieved through debilitation in mesothelial motility since they cannot compete with adhering and proliferating cancer cells for unoccupied serosal spaces.

### Colonization by ovarian cancer spheroids is modulated by the confluence of mesothelial monolayers

To test our computational predictions, we set up coculture time lapses where we parsed between fields where, prior to any prospective colonization event by cancer spheroids (constitutively expressing GFP), the underlying mesothelia were sub-confluent with empty patches and single/dispersed cells lying in between (Figure 6Ai, Bi, and Ci time lapse images: brightfield and GFP at 0 h and 36 h post-acquisition, Supplementary Videos S7-S12), and in another set, confluent (Figure Aii, Bii, and Cii, Supplementary Videos S13-S18). In the sub-confluent mesothelial fields, the spreading ratio for OVCAR-3 spheroids was higher under 50 & 100 µM MG treatment than in control fields (Figure 6D 1.4-fold, 3.4-fold and 4.5-fold (significant) for control, 50 µM, and 100 µM MG respectively; significance computed using one way ANOVA with Tukey’s post hoc comparison). Interestingly though, on confluent mesothelia, even with 50 & 100 µM MG treatments, the invading OVCAR-3 spheroids showed poor adhesion and their spreading ratio was similar to untreated monolayers (Figure 6D 1.3-fold, 1.3-fold and 1.2-fold for control, 50 µM, and 100 µM MG respectively; significance computed using one way ANOVA with Tukey’s post hoc comparison).

**Figure 6.**
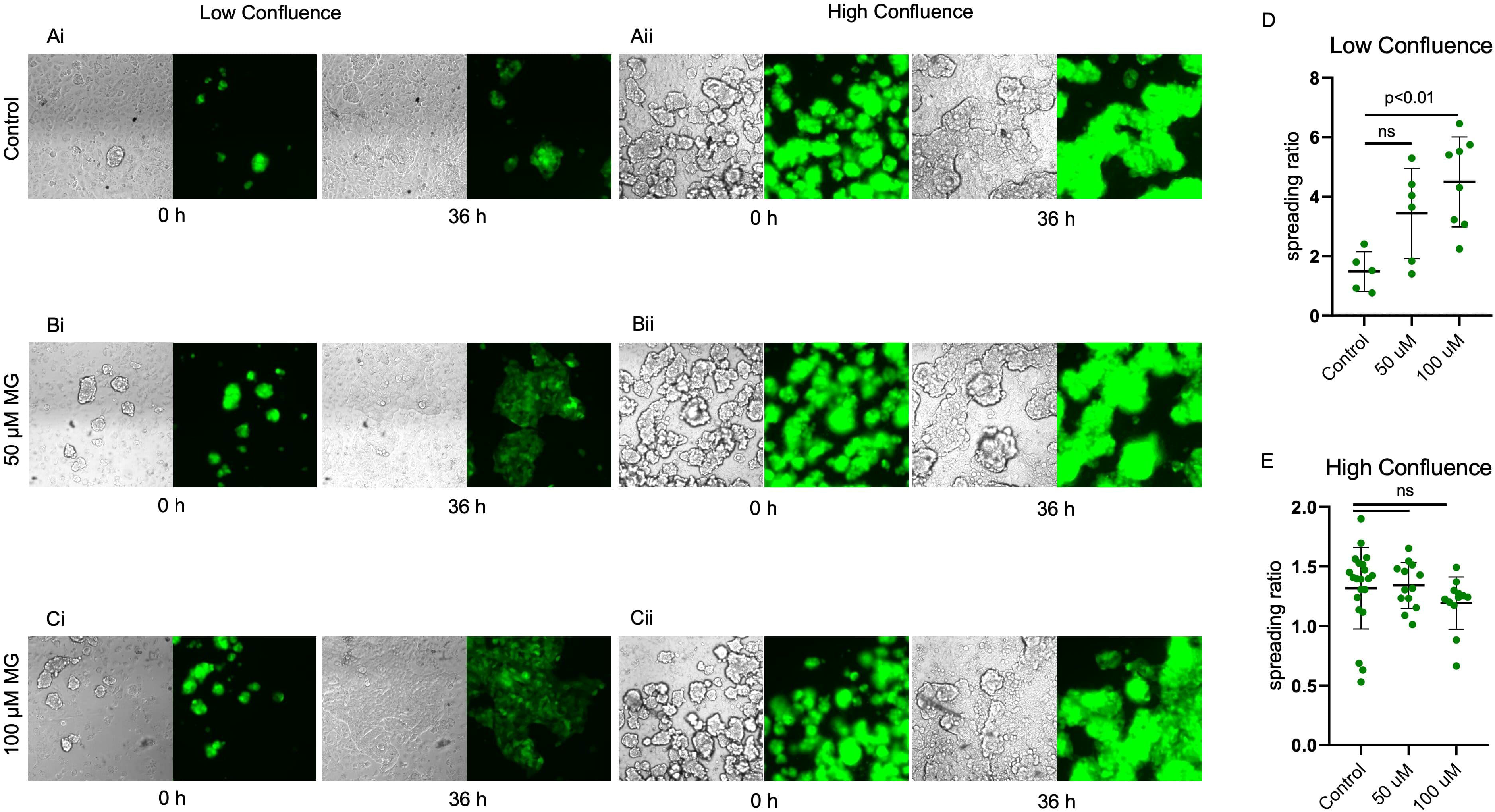
Ovarian cancer spheroid spread more on lower confluence regions of MG-treated mesothelial monolayers. (A) Brightfield and fluorescent micrographs of ovarian cancer spheroids over control and MG-treated untransformed mesothelial monolayers in low (left panel) and high (right panel) confluence at initial (left column) and final (right column) timepoints of cocultures. (B) Graphs plotting the fold change in spreading area in control and MG-treated mesothelial monolayers at low (upper) and high (lower) confluence conditions. Scale bars for A: 100 µM. Quantification done using one way ANOVA with Tukey’s post hoc comparison. Experiments performed n=3 times. (For reference, check Supplementary Videos S7-S18)

### High Glyoxalase-1 levels protect cancer cells from MG

Given that treatment of our cocultures of mesothelia and ovarian cancer cells with MG resulted in a selective toxic effect for mesothelia, we asked if and how cancer cells were protected from the glycation effects of the dicarbonyl compound. To verify this surmise, we treated GFP expressing OVCAR-3 and SK-OV-3 cells with 50 µM and 100 µM MG for 48 h and stained them for propidium iodide to detect dead cells. We observed similar (low) numbers of dead cells with and without MG treatment (Figure 7A). This led us to hypothesize that ovarian cancer cells may express high levels of Glyoxalase-1 (Glo-1), which converts MG to lactate (Nigro et al., 2017). Publicly available data from the Clinical Proteomic Tumor Analysis Consortium (Chen et al., 2023) indicated that ovarian cancer cells had higher levels of GLO-1 than their untransformed controls (Figure 7B) and that GLO-1 levels progressively increased with cancer stages (Figure 7C). We confirmed this using immunocytochemistry and observed higher levels of GLO-1 in SK-OV-3 and OVCAR-3 cells compared with MeT-5A mesothelia (Figure 7D; green: GLO-1 blue: DNA). Viability assays for MG in OVCAR-3 and SK-OV-3 showed IC50 values of 170 and 175 µM, which were much higher than MeT-5A cells (Figure 7E). We then asked if inhibition of Glo-1 in cancer cells can make them more susceptible to MG. We therefore cultured cells with 1 µM, 10 µM, and 20 µM S-p-Bromobenzylglutathione cyclopentyl diester (BBGC), a known GLO-1 inhibitor (Tang et al., 2023) in the absence and presence of 100 µM MG. While we did not observe significant cytotoxicity with the GLO-1 inhibitor alone, in the presence of 20 µM inhibitor, MG decreased the viability of OVCAR-3 cells by more than 50% (Figure 7F; significance computed using one way ANOVA with Tukey’s post hoc comparison).

**Figure 7.**
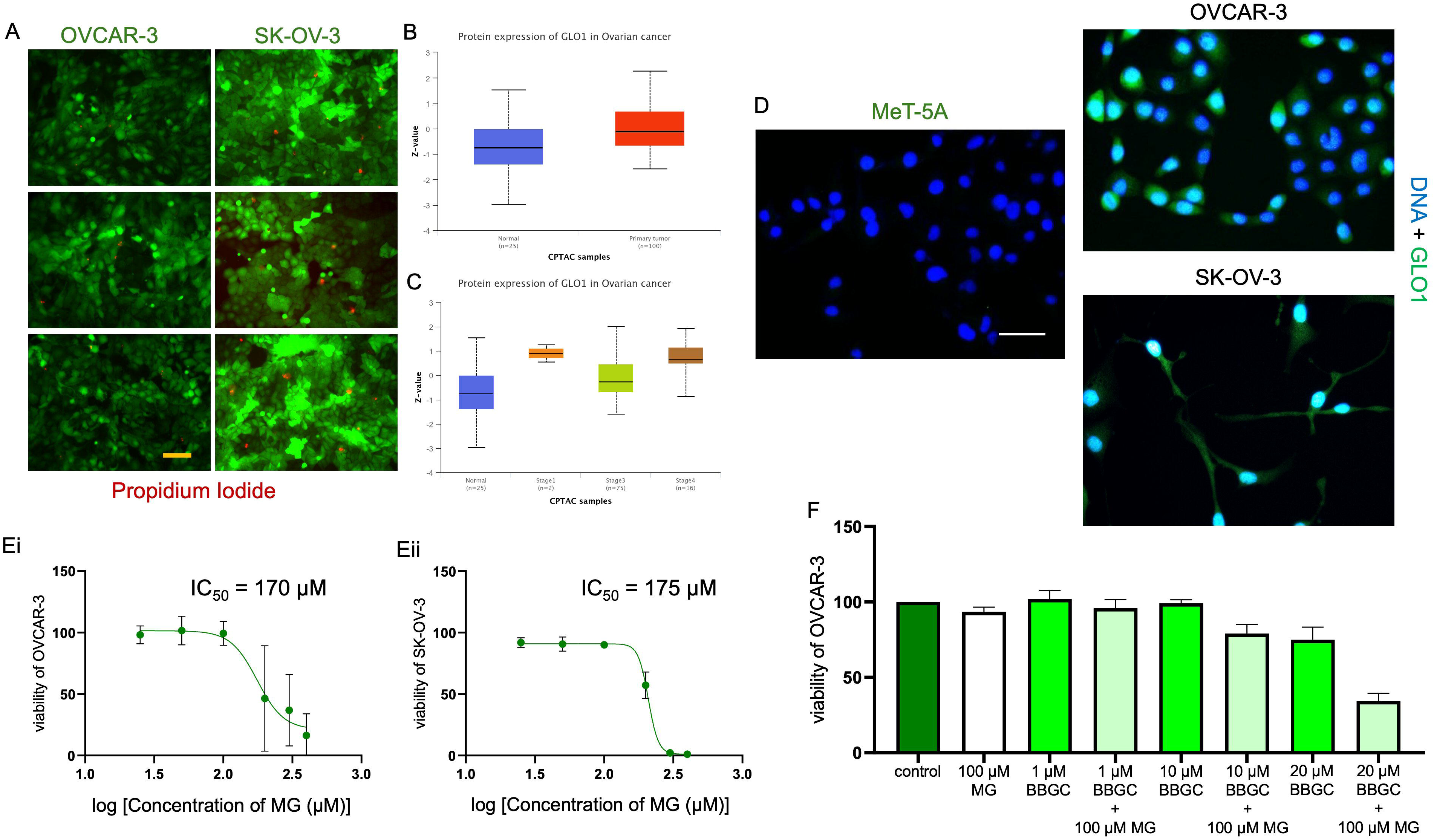
Ovarian cancers are more tolerant to MG due to higher levels of GLO1 expression. (A) Fluorescent micrographs of control and 50 and 100 µM MG-treated OVCAR-3 and SK-OV-3 cells cultured as monolayers and stained with Calcein AM (live; green) and PI (dead; red) markers. (B & C) Protein expression levels of GLO-1 in untransformed ovarian and cancer cells (B) and in stage-stratified ovarian cancer cells (C) as obtained from the UALCAN database. (D) Confocal fluorescent micrographs of MeT-5A, OVCAR-3 and SK-OV-3 stained for DNA using DAPI (blue) and immunostained for GLO1 (green). (E) Graphs plotting viability of (i) OVCAR-3 and (ii) SK-OV-3 in increasing levels of MG. (F) Graph plotting viability of control and MG-treated OVCAR-3 in presence and absence of increasing levels of BBGC (GLO1 inhibitor). Scale bars: 100 µM. Quantification done using one way ANOVA with Tukey’s post hoc comparison. Experiments performed n=3 times.

## Discussion

A seminal paper by Knudson in 1972 established a conceptual bridge between hereditary and non-hereditary forms of cancer by postulating that familial malignancies are driven by one inherited germ-line mutation and another somatic mutation (Knudson, 1971). In a recent paper on pancreatic ductal carcinogenesis, Kong and coworkers build on this two-hit hypothesis by showing that for BRCA2, even if one allele is mutated, tumorigenesis can be rapidly initiated by the excess production of a reducing glycolytic metabolite: methylglyoxal (MG) which brings about the proteolysis of the protein translated from the other BRCA2 allele (Kong et al., 2024) and therefore manifests its haploinsufficiency (Jiang, 2024)). Cancer cells frequently reprogram their metabolic dynamics to upregulate glycolytic flux primarily to generate building blocks for anabolic biomolecular synthesis as well as to generate energy in combination with an upregulated oxidative phosphorylation (Chandel, 2014; DeBerardinis & Chandel, 2024). Therefore, an increased generation of MG, a glycolytic byproduct is inevitable under such switched onco-metabolic contexts (Rabbani et al., 2016).

However, for a transformed tumor to spread through the body, multiple suppressive mechanisms along the trajectory of spread need to be in turn suppressed. Research over the last few decades strongly attributes the milieu surrounding the once-transformed tumor cell(s): the extracellular matrix and the stromal cells, as demonstrating complex roles in regulating the kinetics of tumor progression. The effect of this milieu: the tissue microenvironment is in turn strongly regulated by macroenvironmental-systemic cues (Bhat & Bissell, 2014). A case in point, Type 2 Diabetes mellitus (T2DM) is considered a risk factor for cancers such as hepatocellular carcinoma (Negro, 2020). Herein, the same advanced glycation end products that are formed by interactions of MG with proteins tend to alter the rheological properties of the extracellular matrix such as their viscoelasticity (Fan et al., 2024). This exacerbates the progression of hepatocellular carcinoma in a integrin-β1–tensin-1–YAP-dependent manner. In this paper we sought to ask how the accumulation of AGE through MG may alter the serosal homoeostasis that is established by stromal cells against an invasive onslaught of colonizing cancer spheroids. Following up on these lines, we were also motivated by recent work that shows how untransformed epithelial cells can extrude transformed cells within their population and maintain quiescence within multicellular sheets (Vishwakarma & Piddini, 2020).

A corollary of our study is that healthy confluent mesothelial monolayers are able to impair the attachment and spreading of cancer spheroids. This can be deduced based on our experiments and simulations on their ability to tightly appose with each other using intercellular junctions but also fill up unoccupied substrata through proliferation and motility. Mesothelia are unique in their ability to quickly move to denuded peritoneal surfaces both by surface locomotion as well as by shedding and re-adhesion (Yung & Davies, 1998). In addition, the tight apposition of epithelia results in a mesoscopic scale elastic behavior through synchronization of actin cytoskeleta by proteins that bind to it and link it with membrane proteins especially on the apical surfaces such as Ezrin and ZO-1 (Doctor, 2006; Van Citters et al., 2006). The dual property of apposition and motility of mesothelia, seems to us crucial to its ability to prevent cancer spheroids from adhering and spreading. Impairment in both these properties can facilitate ovarian cancer spheroids that have attached on acellular spaces to disintegrate and spread through cell division and migration. On the other hand, interspersion of acellular patches within a mesothelial monolayer can also weaken the global elastic properties of the layer rendering it susceptible to spheroidal adhesion and clearance even in the confluent zones. Future investigations will test the spatial congruence of the extent of confluence and the initial attachment sites of spheroids.

The impairment of their localization by MG could both be a direct or indirect effect of protein glycation and will be investigated in greater details in subsequent investigations. Glycation affects not just proteins but nucleic acids as well (Prasad et al., 2019), and therefore the effects of MG on mesothelia could be driven by (epi)genetic alterations leading to a qualitatively distinct mesothelial transcriptome and proteomes. We aim to undertake omic investigations in follow up papers to identify the mechanisms of disruptions in localization of key cytoskeletal regulators seen in our study.

It has not escaped our notice that ovarian cancer cells have elevated levels of GLO-1 which doesn’t just detoxify MG but in fact converts it to lactate (Distler & Palmer, 2012; Yang et al., 2022). Lactate is considered a putative oncometabolite that is used by cancer cells to acidify its microenvironment, resulting in immunosuppression and angiogenesis, and ultimately tumor progression (Gillies et al., 2019; Pérez-Tomás & Pérez-Guillén, 2020; San-Millán et al., 2020). Therefore, the very dicarbonyl may act as a fuel for the cancer cells and a poison against mesothelial defense against cancer. Our results with Glo-1 seems to restore susceptibility of cancer cells to MG suggesting that future precision therapy investigations should examine the potential for combination of Glyoxalase inhibitors with chemotherapeutics for patients with high levels of glycation caused by T2DM or simply aging.

Finally, in this study, the effect of T2DM-mimicking microenvironment was achieved through treatment using only a single dicarbonyl, MG, whereas other glycating agents like 3-deoxyglucasone and glyoxal are known to be elevated in diabetes and contribute to the formation of AGEs such as pentosidine and pyrraline (Ashraf et al., 2015, 2016; Niwa, 1999). In future studies we will extend our investigations to diverse dicarbonyls as well as to other stromal tissues to develop a more integrated histopathological theory on how selective stromal glycation potentiates ovarian cancer metastasis in the peritoneum.

## Acknowledgements

This work was supported by the Wellcome Trust/DBT India Alliance Fellowship/Grant [IA/I/17/2/503312] awarded to RB. It was also supported by the John Templeton Foundation (#62220) and the Indo-French Centre for the Promotion of Advanced Research (69T08-2) to RB. SM (Satyarthi) and SM (Sudiksha) acknowledge their Prime Ministers Research Fellowship. CVSP and AG acknowledge the Axis Bank Ph.D. Fellowship and support from the Ministry of Education, Government of India, respectively. The opinions expressed in this paper are those of the authors and not those of the John Templeton Foundation.

